# Characterising maize viruses associated with maize lethal necrosis symptoms in sub Saharan Africa

**DOI:** 10.1101/161489

**Authors:** I.P. Adams, L.A. Braidwood, F. Stomeo, N. Phiri, B. Uwumukiza, B. Feyissa, G. Mahuku, A. Wangi, J. Smith, R. Mumford, N. Boonham

## Abstract

Maize lethal necrosis disease (MLN) is an emerging disease in East Africa caused by the introduction of *Maize chlorotic mottle virus* (MCMV). Recent activity seeking to limit spread of the disease is reliant on effective diagnostics. Traditional diagnostics applied on samples with typical field symptoms of MLN have often given negative results using ELISA or PCR for MCMV and *Sugarcane mosaic virus* (SCMV). Samples collected in the field with typical MLN symptoms were examined using next generation sequencing (NGS). SCMV was found to be more prevalent than suggested by targeted diagnostics. Additionally, the panel of samples were found to be infected with a range of other viruses, seven of which are described here for the first time. Although not previously identified in the region, *Maize yellow mosaic virus* (MYMV) was the most prevalent virus after MCMV. The development of targeted diagnostics for emerging viruses is complicated when the extent of field variation is unknown, something that can be negated by using NGS methods. As a result we explored MinION technology which may be more readily deployable in resource poor settings. The results show that this sequencer can diagnose known viruses and future iterations have the potential to identify novel viruses.

## Introduction

Maize is a major commercial and subsistence crop in sub - Saharan Africa accounting for 34% of all cereals grown ^1^. Maize Lethal Necrosis (MLN) is a serious viral disease affecting maize production. First identified in the USA in 1976 ^2^ the disease had largely remained within the Americas (Peru, Argentia, Mexico). First reports have recently been recorded for China in 2011 ^3^and Africa (Kenya) in 2012 ^4, 5^. Since the initial outbreak in Kenya, MLN has now been reported in Tanzania^6^, Rwanda ^7^, Ethiopia ^8^, the Democratic Republic of Congo ^9^ and South Sudan ^1^, and is likely to be of wider distribution.

Maize lethal necrosis (MLN) is caused by a co - infection between *Maize chlorotic mottle virus (*MCMV) and a potyvirus, in recent outbreaks this is usually *Sugarcane mosaic virus (*SCMV). Other members of the potyviridae, including *Maize dwarf mosaic virus* ^10^ or *Wheat streak mosaic virus*^6^, have also been associated with the disease. It is thought that the second virus supresses the host plants post - transcriptional gene silencing based antiviral defences allowing for increased replication of MCMV and subsequent devastating disease ^6^. Field surveys using ELISA in Kenya in 2013 - 14 showed that 60 % of randomly selected samples were positive for MCMV, whilst only 28% were infected with SCMV ^8^. However, interpretation of this data is complicated by a poor understanding of the nucleic acid sequence variability within the virus population of the region and the impact (efficacy) this may have on the diagnostic assays used. MCMV within sub Saharan Africa has been found to be highly conserved ^8^, and as a result nucleic acid based assays for MCMV should be reliable. However, the commercially available antibodies used in the ELISA testing were not raised to isolates from the region and have subsequently been found to give negative results with some infected samples from Kenya ^1,5^. *Sugarcane mosaic virus*, has much higher nucleotide sequence diversity,^7^ and a real - time PCR assay designed to the sequence of the original Kenyan isolate ^5^ was found not to amplify SCMV isolates from Rwanda ^7^. There is also evidence of uncertainty when trying to identify maize lethal necrosis using field symptoms alone. Maize samples obtained from Democratic Republic of Congo with severe chlorotic mottle and pale green streaks typical of MLN were found not to be infected with a potyvirus (using either SCMV specific or generic potyvirus primers). *Maize streak virus* was also found to be absent ^9^. It seems likely that disease symptoms caused by other well characterised viruses are being confused with MLN in the field or that different potyviruses may be contributing to the disease in some instances and that other, as yet undescribed viruses may also be contributing to symptoms and subsequently confusion around identifying maize lethal necrosis accurately in the field.

Next generation sequencing (NGS) technology, first used for plant virus detection in 2009 ^11-13^ has since been used a number of times to identify the cause of crop diseases ^14^. However, such technology is largely not available in Africa due to high cost of equipment and service support and reagent costs. The introduction of simpler, table top sequencers such as the MiSeq (Illumina) and Ion Torrent (Life Technologies) has made this technology easier to access, bringing down both the capital cost of the equipment and per sample cost of the reagents. Despite this, the technology is still only available in specialist centralised diagnostic laboratories ^15^. The current technology relies on detecting the incorporation of nucleotides into a second DNA strand complimentary to the target DNA in a process collectively described as sequencing by synthesis (i.e. 454, Ion Torrent, Illumina MiSeq / HiSeq / GAII, Pacific Bioscience RSII), or sequencing by ligation (i.e. Life technologies SoLiD). The detection of these reactions requires a sensitive instrument which contributes to the expense of the systems. For over 10 years the potential of sequencing DNA by passing it through a nanopore has been discussed ^16^. This approach was thought to offer potential benefits in terms of equipment simplicity and hence price, the length of reads generated by the sequencing pores and the pre - processing required for the sample before it could be sequenced. However, it is only recently that a commercial device based on this concept (MinION, Oxford Nanopore Technology Ltd.) has become available ^17^. The MinION sequencer has reduced the cost of next generation sequencing instruments by 50x, in addition since it relies on remote, cloud based analysis there is not a requirement for powerful on - site computing platforms, thus it may be useful for implementation in resource poor settings such as Sub Saharan Africa. The MinION has recently been used for the sequencing of viral cDNA amplicons during the 2015 Ebola outbreak ^18^ as well as for the detection and characterisation of isolates of Chikungunya, Ebola and Hepatitis C ^19^ and poxviruses from human blood ^20^.

The current study used ‘standard’ MiSeq and HiSeq sequencing, to investigate viruses that were present in samples from several countries where field symptoms of MLN were reported, but the presence of MCMV and SCMV could not always be confirmed using laboratory testing. In addition a comparative analysis was completed using the MinION sequencer, investigating its effectiveness for generating viral genome sequences enabling the identification of both new and previously characterised viruses.

## Results

Leaf samples (n = 66) of maize from 26 sites in four countries, all identified with typical field symptoms of MLN disease, were collected. For each site the samples were pooled and tested using real - time PCR or ELISA (targeted diagnostics) and NGS. The results (Table 1) show that MCMV was detected at 25/26 of the sites using targeted diagnostics and NGS, whilst SCMV was found at 18/26 and 22/26 sites using targeted diagnostics and NGS respectively. SCMV was only found in combination with MCMV.

**Table 1.**
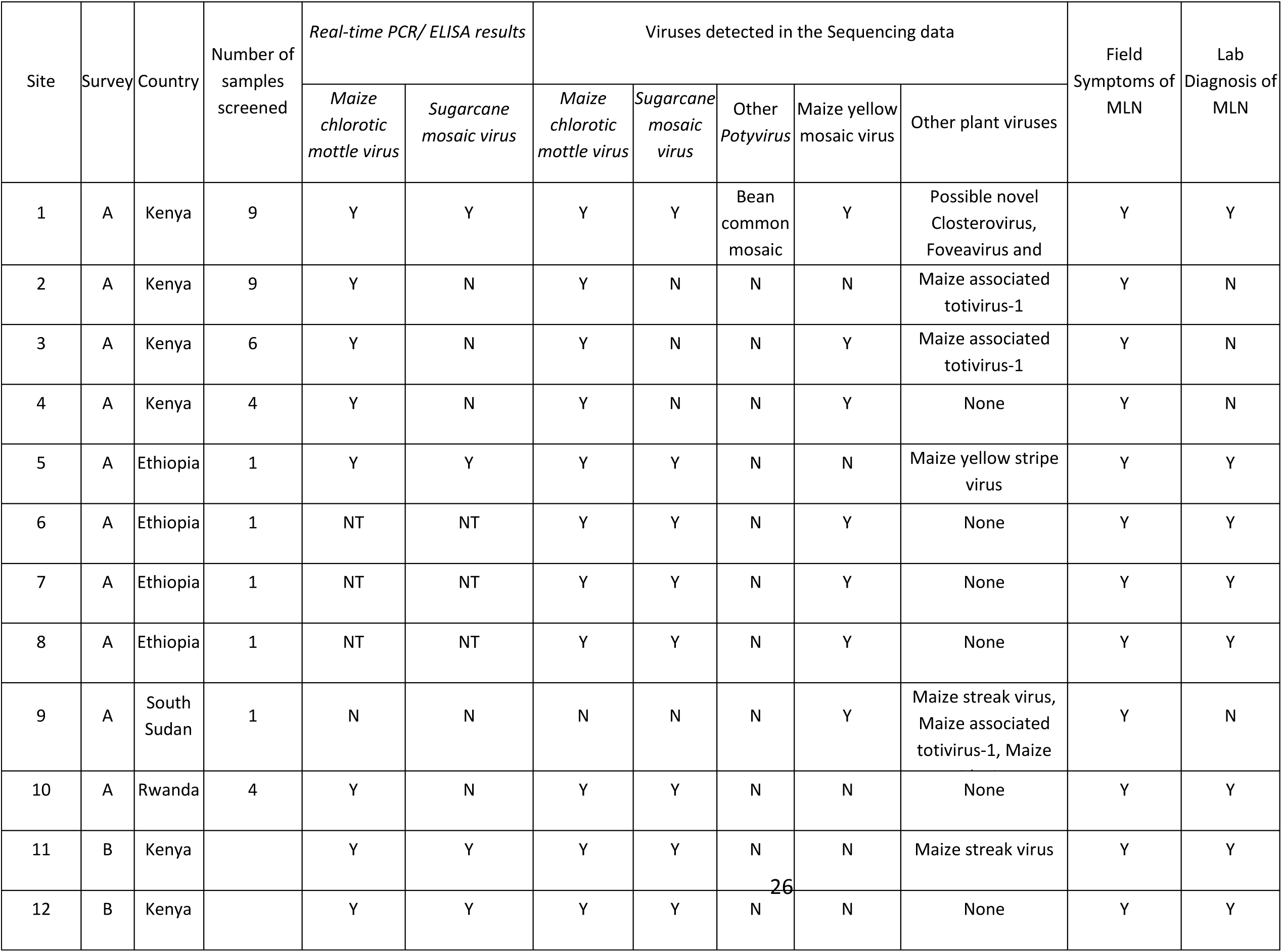
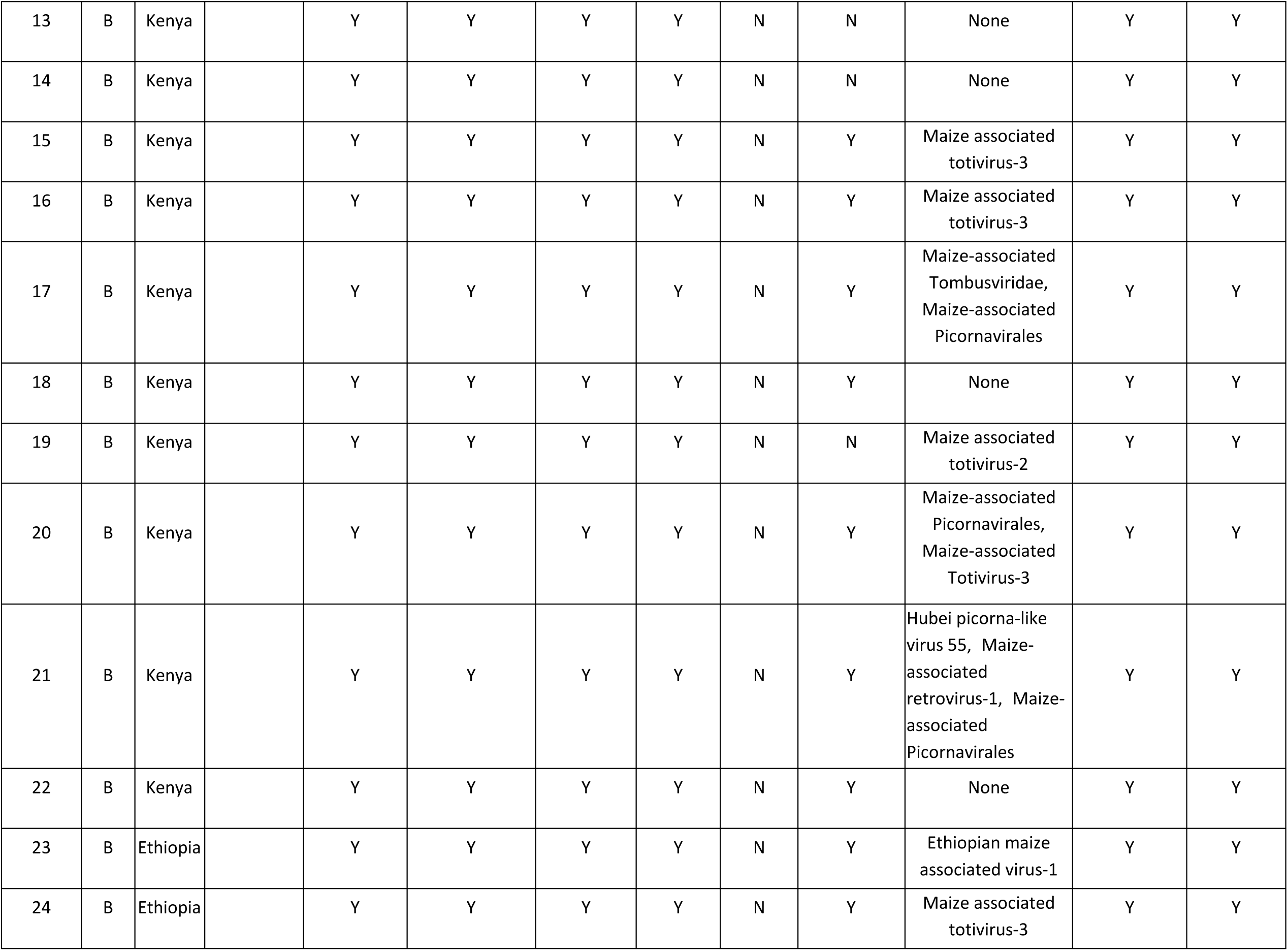
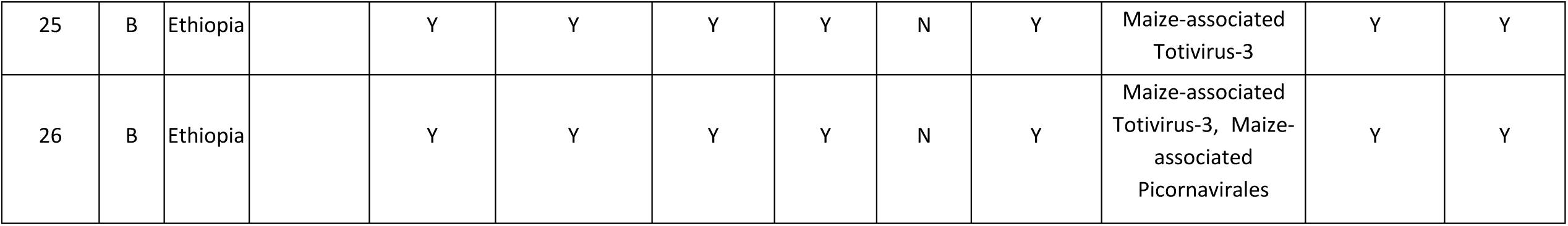
Details of sites investigated, including symptoms observed, targeted diagnotics results and the virus sequences identified in the NGS data for each of the samples tested.

A number of other previously described viruses were also recorded. *Bean common mosaic virus, Maize yellow stripe virus* (MYMV), *Maize streak virus* (MSV) were each recorded in one site and *Maize yellow mosaic virus* (MYMV) was recorded in 19 sites. Novel sequence motifs were also identified with similarity to members of the *Closterovirus, Foveavirus, Amplelovirus, Totivirus, Partitivirus* and *Pteridovirus* genera, Tombusvirdae family and Picornavirales order. Namely: A 1,059nt contig (acc MF372909) was identified from one site encoding a putative ORF with a *Closterovirus* coat protein motif ^21^ and 32% amino acid identity to the minor coat protein of Coryline virus 3 (acc AGF73885).

A 3,009nt contig (acc MF372910) was identified from one site encoding for two ORFs with viral helicase and RdRp motifs and similarity to members of the *Foveavirus* genera. The contig had 64% identity to the replicase and 41 % identity to the triple block protein of Asian prunus virus 2 (acc KR998048) an unclassified *Foveavirus*.

A 1,288nt contig (acc MF372911) was identified from one site encoding an open reading frame for a protein with 36% amino acid identity to the methyltransferase of Grapevine leafroll associated virus 3, from the genus *Amplelovirus*.

A 2985nt contig (MF415880) was identified from one site encoding an open - reading frame with 66% nucleotide identity to *Citrus yellow vein - associated virus*, an unclassified virus, and a conserved viral RdRP 3 superfamily domain. It was designated as *Ethiopian maize - associated virus - 1*.

A 9075nt contig (MF415876) was identified from one site, containing conserved domains associated with single stranded RNA viruses: two picornavirus capsid domains and a P - loop. It also contains a cricket paralysis virus capsid domain, and a reverse transcriptase domain. This may represent a novel retrovirus or mobile genetic element, and was designated as *Maize - associated retrovirus - 1*.

One site contained *Maize - associated totivirus* 2 (MTV - 2) a 4427nt (MF415877) contig with 99% nucleotide identity to the first reported sequence from Ecuador (KT722800). Six sites contained contigs with 81% nucleotide ID to KT722800, which may represent a divergent strain of MTV - 2 or a novel Totiviridae species. These sequences were designated as *Maize - associated totivirus - 3* (Acc:MF425844 - 9). One sample contained a 9869nt contig (Acc MF415875) with 99% nucleotide identity with *Hubei picorna - like virus 55*, recently discovered in a mixed sample of Chinese spiders^22^. This is its first report outside of China.

Four sites contained contigs which had discontinuous megablast hits to *Soybean associated bicistronic virus* (KM015260) and *Aphis glycines virus 1* (KF360262), newly discovered members of the Picornavirales which appear to be the same virus (99% nucleotide ID between them). The contigs had homology covering 67% with 75% nucleotide identity, suggesting that this represents a novel Picornavirales member, designated as *Maize - associated picornavirales (*MF425850 - 55).

A 4,221nt (MF415879) contig was identified which has a discontinuous megablast hit to *Maize white line mosaic virus* (Aureusvirus: Tombusviridae), with 69% nucleotide ID over 20% of the contig. Reading frame analysis revealed ORFs consistent with the Tombusviridae, including the characteristic read - through expression of the RNA - dependent RNA polymerase (RdRP). Combined with conserved domain search results consistent with the Tombusviridae family, this likely represents a novel Tombusviridae member (designated as *Maize - associated tombusviridae*) in a new genus - given its divergence from previously discovered members of the family.

Two contigs 5,741nt and 2,693nt were identified that shared 43% and 31% protein identity with parts of the RNA1 and RNA2 encoded polyproteins from Japanese holly fern mottle virus (JHFMV) with which they appear to share the same genome organisation (Valverde & Sabanadzovic, 2009). The 5741nt RNA1 molecule (acc MF372912) encodes a putative single 2,013kDa polyprotein with methyl transferase, viral helicase and RdRp motifs (Marchler - Bauer et al., 2015). The 2,693nt RNA2 molecule (acc MF372913) putatively encode 3 proteins the first contains a movement protein motif with 33% identity to the JHFMV movement protein and the third showing 23% identity to the JHFMV coat protein. The middle open reading frame encodes a putative 30kDa protein with no significant homology to any sequences on Genbank and may be analogous to the JHFMV p37 although it has little amino acid identity. This virus may be the second member of the proposed genus *Pteridovirus* and we suggest the name Maize pteridovirus - 1 (MPtV - 1).

Four distinct isolates of Maize associated totivirus 1 (MTV - 1) ^23^ were also identified. Two contigs of lengths 4,987nt and 3,065nt (acc MF372914, MF372915) had 86% and 67% identity respectively to MTV - 1. One contig of 4,867nt (acc MF372916) had 85% identity and one contig of 4,986nt (acc MF372917) had 81% identity MTV - 1.

A 1,849 nt (acc MF372918) contig was identified that appears to encoded a putative ORF of 67 kDa protein containing a RNA - dependent RNA polymerase motif ^24^. The contig has 34% identity to a 66kDa protein encoded by *Penicillium aurantiogriseum* partiti - like virus ^25^. For the purposes of this study this putative virus was called Maize associated partiti - like virus (MAPLV).

Of the sites studied, 69% also had the recently discovered Maize yellow mosaic virus (MYMV). Figure 1 shows a neighbour joining tree produced from coat proteins of the MYMV sequences available on GenBank and produced from this study. The sub - Saharan African isolates of MYMV form a distinct cluster (<99% similarity), but are similar (98 - 99%) to Chinese isolates.

**Figure 1.**
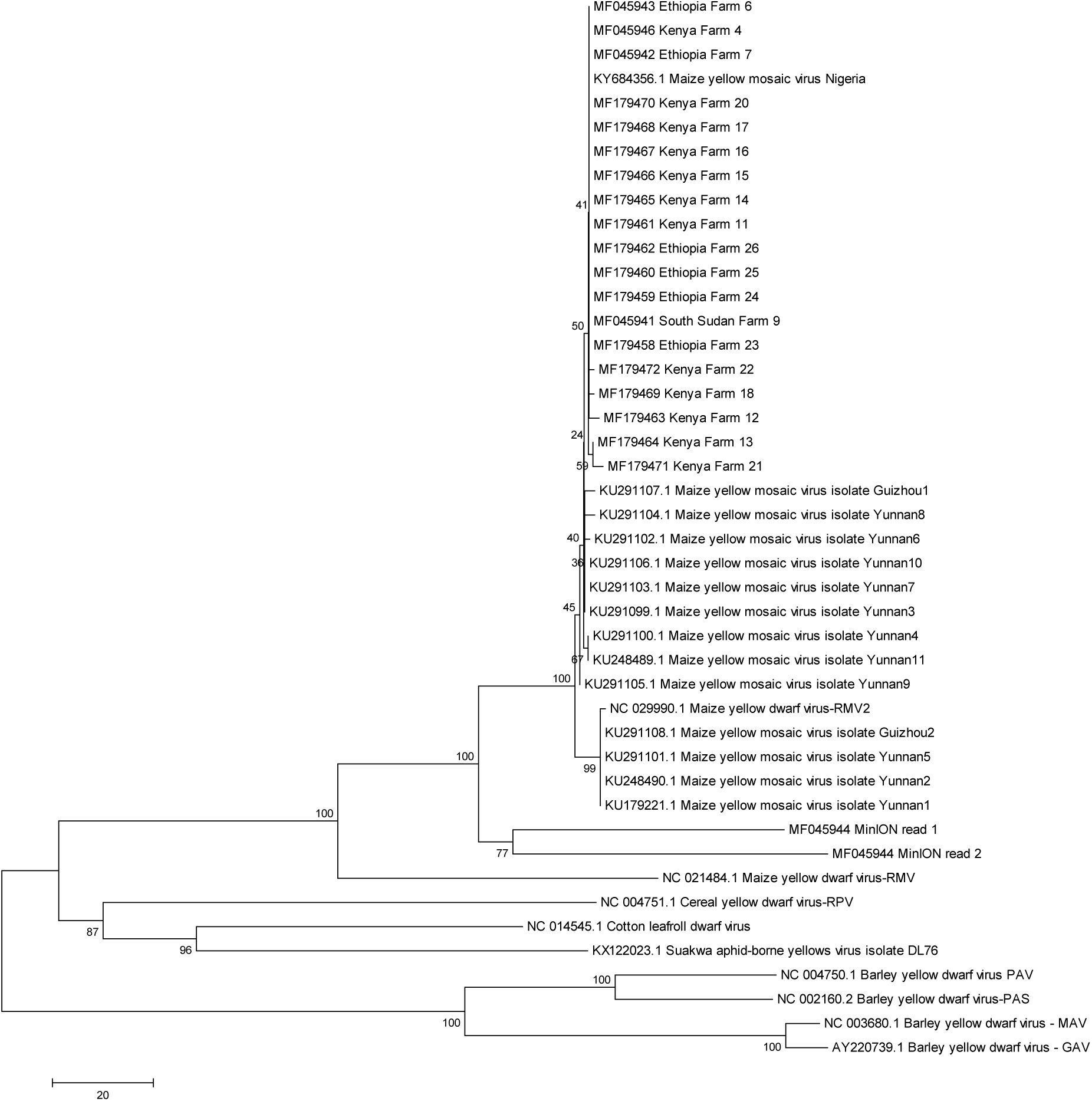
Maximum likelyhood tree of Maize yellow mosaic virus coat protein sequences and related virus coat proteins. The sequences generated using the MinIon platform are included in the analysis.

RNA from samples of a single site from South Sudan was sequenced on a single run of an R7.3 MinION flow cell. MiSeq sequencing of the same sample had yielded 768,902 read pairs and BLASTn analysis using the October 2015 Genbank database had shown the presence of MSV with 99% nucleotide identity to an isolate from Mozambique (Acc: FJ882098) ^26^ and a virus with 81% nucleotide identity to the sequence of the recently reclassified *Maize yellow dwarf virus* ^27^. Further examination using a more recent BLAST database revealed this virus to be MYMV ^28^. BLASTx analysis revealed the presence of MPtV - 1, MAPLV and two distinct isolates of MTV - 1 (described above).

Over a 48hr period the MinION produced a total of 109,560 reads from 331 pores, yielding 9,560 high quality 2D “pass” reads. The longest was 8,045nt and the average read length was 1,130 ± 541. (SRA acc: SUB2651212) (Figure 2a). Of the reads produced, approx. 50% were during the first 2 hours and 75% in the first 7 hours (Figure 2b). A further 38,784 lower quality, 2D reads were recovered from the “fail” output with an average length of 895 ± 498. The high quality reads were compared to the GenBank nr and nt databases from October 2015. Following BLASTn, sequences with significant identity to two viruses were identified. A total of 24 reads (between 609 and 3,857nt in length) had up to 87% identity to sequences of *MSV* and 3 reads (3,188nt, 2,100nt & 1,284n) had up to 82% sequence identity to *Barley yellow dwarf virus* (subsequently identified as MYMV). Following BLASTx analysis, sequences with significant homology to these viruses were again found with the addition of sequences with significant identity to MTV - 1. No sequences with significant sequence identity to either MPtV - 1 or MAPLV were found.

**Figure 2.**
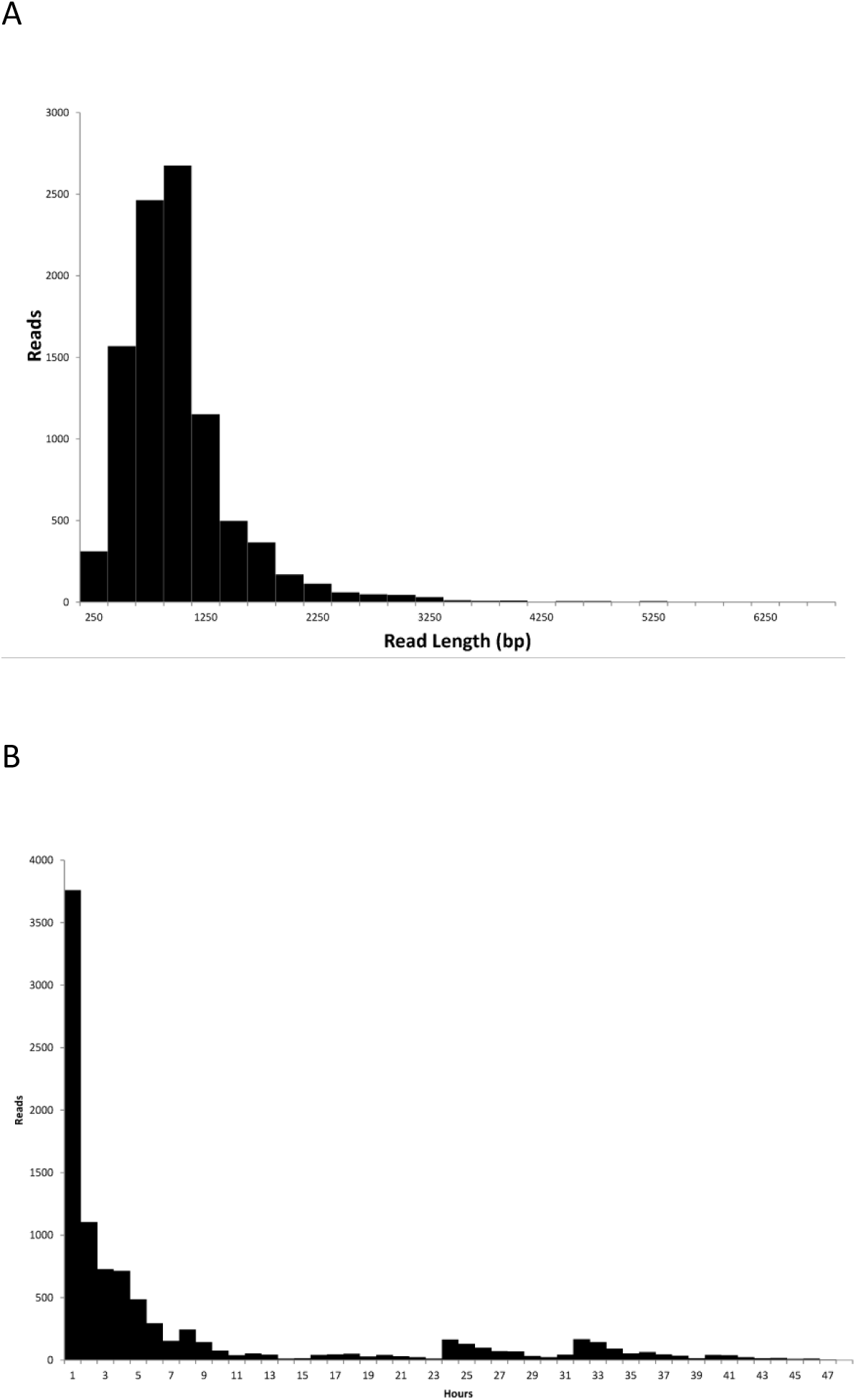
(A) Histogram showing size distribution of high quality MinION reads. (B) Histogram showing time when high quality MinION reads were produced.

In order to determine if sequences corresponding to MPtV - 1 and MAPLV were generated during the MinIon run all the high quality sequencing reads were mapped on to the assembled viral genomes identified using the MiSeq data. All 5 viruses were present in both sequencing datasets (Table 2) in approximately similar proportions.

**Table 2.**
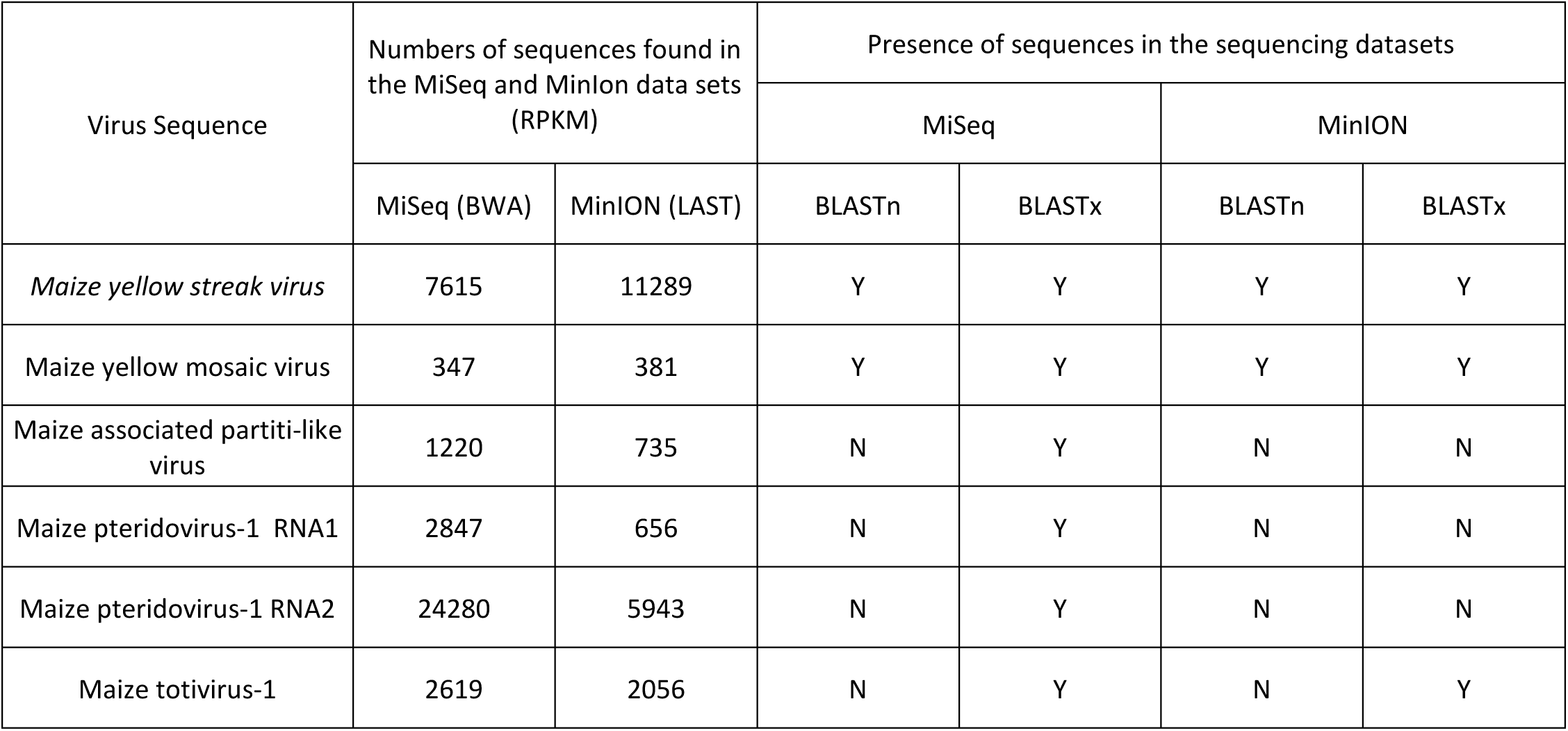
Comparison of the presence (Y) or absence (N) and quantity of virus sequences identified in the MiSeq and MinION data for the South Sudan sample.

| Virus Sequence | Numbers of sequences found in the MiSeq and Minlon data sets (RPKM) |  | Presence of sequences in the sequencing datasets |  |  |  |
| --- | --- | --- | --- | --- | --- | --- |
|  |  |  | MiSeq |  | MinION |  |
|  | MiSeq (BWA) | MinION (LAST) | BLASTn | BLASTx | BLASTn | BLASTx |
| *Maize yellow streak virus* | 7615 | 11289 | Y | Y | Y | Y |
| Maize yellow mosaic virus | 347 | 381 | Y | Y | Y | Y |
| Maize associated partiti-like virus | 1220 | 735 | N | Y | N | N |
| Maize pteridovirus-1 RNA1 | 2847 | 656 | N | Y | N | N |
| Maize pteridovirus-1 RNA2 | 24280 | 5943 | N | Y | N | N |
| Maize totivirus-1 | 2619 | 2056 | N | Y | N | Y |

The MinION sequence data for MYMV was compared with that generated using the MiSeq and the sequences from Genbank. The results (Figure 1) show that the MinION reads clustered with the sequences generated using the MiSeq, in a cluster within the MYMV sequences, distinct from other related *Polerovirus* and *Leutovirus* sequences.

The MinION reads were uploaded to the WIMP tool. Within 20 minutes of the WIMP analysis being started, taxonomic calls were being returned for the initial sequences. The output from the 109,560 MinIOn reads (Figure 3) showed that WIMP was able to correctly identify the presence of MSV and MYMV. Sequences of MPtV - 1, MTVs or MAPLV were not identified as having homology to virus sequences. Sequences were found with a low - confidence score (<0.0106) to *Choristoneura occidentalis granulovirus,* a dsDNA *Betabaculovirus* infecting western spruce budworm. Sequence data could not be recovered using the WIMP package and thus it was not possible to evaluate this sequence further.

**Figure 3.**
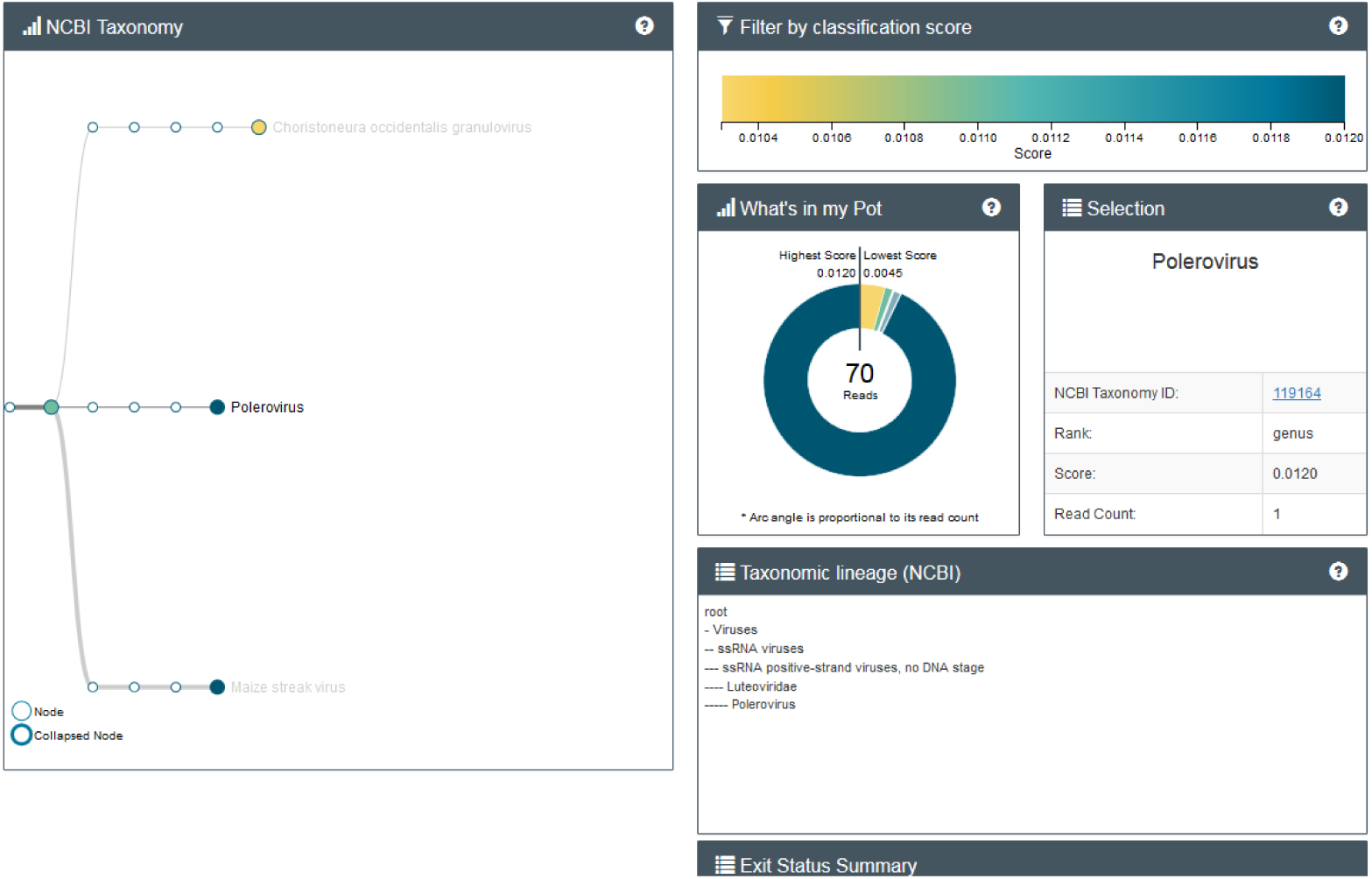
Output from WIMP tool.

## Discussion

Maize is an important commercial and subsistence crop in sub Saharan Africa and production is under threat from the emergence and rapid spread of the viral disease Maize Lethal Necrosis (MLN) ^1^. This study used targeted (real - time PCR / DAS - ELISA) and non - targeted nucleotide sequencing to examine maize leaf samples with symptoms of MLN from 26 sites in 4 countries. In survey A, MCMV was detected in samples from 9 sites by both methods, however, SCMV was only detected in 2 sites using real - time PCR and 6 sites using sequencing. Thus, a virus complex characteristic of causing MLN was only confirmed for 6 of the 10 samples identified with MLN based on field symptoms. Of the remaining four sites which had symptoms of MLN, two sites had MCMV and MYMV, one site had MSV and MYMV and one just MCMV. In survey B, all samples were positive for MCMV and SCMV, both by DAS - ELISA and NGS. Using a non - targeted sequencing approach we were able to identify other viruses present in the samples from all of the sites. MYMV was found at 19 of the 26 sites across both surveys, 16 in combination with MCMV and SCMV, two were in combination with MCMV and one had MYMV and MSV.

MYMV was first described and named in China in 2016 ^29^, followed by the sequencing of the whole genome ^28^ and has recently been reported in Nigeria ^30^. The symptoms are described as “yellow mosaic” ^28^ or “yellowing and dwarfing” ^29^. The symptoms shown in Chen et al. ^28^ appear at least superficially to be similar to those of MLN. This study is the first report of MYMV in sub Saharan Africa and the second report outside China. Its occurrence in multiple sites in 3 different countries in sub - Saharan Africa suggests that it may be wide spread. The sequences of the isolates of MYMV from Africa were all similar to each other and distinct to the published isolates of China. MYMV is a polerovirus which, like potyviruses, are known to suppress post transcriptional gene silencing ^31^, although using a different mechanism; it would be interesting to investigate if MYMV, damaging in its own right is also capable of enhancing the damage caused by MCMV.

In addition to the characterised viruses found in the samples, 7 novel virus - like sequences were detected. These include potentially novel species from the *Closterovirus*, *Amplelovirus*, *Foveavirus, Partivirus* and *Pteridovirus* genera, and two further potentially novel species that could not be classified below the family level – Picornavirales and Tombusviridae. It is not clear if any of these viruses are associated with symptom development in the plants tested.

This project highlights the issues with rapidly introducing targeted diagnostics for emerging virus problems, where the variability of the target is not yet fully elucidated. For MLN diagnostics this has impacted both the ELISA and PCR testing for MCMV and SCMV, respectively. Non - targeted diagnostics (i.e. next generation sequencing) on the other hand circumvent these issues, however the technologies are typically only found in large centralised testing facilities due to the capital costs of the technologies being used and the complexity and infrastructure needed for the analysis. The introduction of the MinION (Oxford Nanopore) opens the possibility of sequencing technologies being available in resource poor, extension and regional diagnostics labs where currently NGS analysis is unavailable. This potential has been demonstrated by the use of nanopore sequencing in the recent Ebola ^18^ and Zika outbreaks ^32^.

This study examined the potential for MinION sequencing to be used for identifying plant viruses in diseased plant material. Using the sequencing data generated using the MinION technology we were able to identify the presence of *Maize streak virus, Maize yellow mosaic virus* and Maize totivirus 1. However, by comparison to the MiSeq data the high error rate of the MinION (approximately 13% including many indels which prevented accurate amino acid sequence determination stopped the identification of 2 novel viruses. Reference mapping MinION sequences to viral genomes did however confirm that sequences for each of these viruses were present in the data set.

Based on these observations, using the R7 flow cells could be useful as a screening tool for testing samples for the presence of well characterised viruses, but may not be suitable for detection of divergent targets or unknowns. The release of the R9 flow cells (with a predicted 5 - 10% error rate) may overcome this situation and provide a level of analysis more similar to MiSeq sequencing and, therefore, virus discovery applications. Whilst the MinION sequencing device itself was simple to use and would be suited to routine testing applications, the sample processing required before the sample is applied to the flow cell is similarly time consuming and complex as other sequencing platforms. This may limit the uptake of the MinION into routine testing laboratories until the protocol can be streamlined, or the potential of MinION based direct RNA sequencing is realised. In addition to the sequencing platform we explored the use of the WIMP application for data analysis. Whilst the system was not as effective as conventional bioinformatics approaches for identifying novel virus sequences, it was very simple to use and well suited to use by people not experienced in bio - informatics. In addition, the use of cloud computing removed the need for powerful computer servers for analysis, something that currently reduces the potential uptake of NGS techniques into routine diagnostic laboratories.

MLN is a damaging disease of maize crops and is currently spreading throughout sub Saharan Africa. Currently surveillance for the disease is predominantly based on observations of the disease symptoms. Our study demonstrates that not all samples with MLN - like symptoms are caused by the disease and that a number of new, or recently discovered viruses, are likely to also be associated with these symptoms. Whilst it is impossible from the current work to identify which viruses are associated with symptoms in maize, it does highlight the need for the deployment of specific diagnostics for the causal agents of MLN disease. The results also suggest that a greater knowledge is needed of the additional viruses and the risk these may be presenting with production and with spread through the seed industry. It is likely that certification of maize seed cannot be effectively based on visual observations and that a number of specific and sensitive diagnostics are required to measure for a number of known high - impact viruses along with viruses that may be of potential harm. Non - targeted sequencing methods are useful for identifying these other viruses, however due to the costs and infrastructure needs of the most widely used, existing platforms, uptake is slow. The applicability of such technology to certification schemes is further challenged by virtue of the numbers of samples that may need to be tested. Nevertheless, the MinION system examined here shows promise for both virus discovery and for identification of the virus variability, and both the costs of the platform and the nature of the analysis offer benefits for studying emergent diseases in resource - poor settings.

## Experimental Methods

### Samples

Survey A: Samples were collected on farms from the Bomet (2012) (site 1 and 2) and Trans Nzoia (2013) (sites 3 and 4) regions of Kenya, Ethiopia in 2014 (sites 5, 6, 7, 8) as described in ^8^, in south Sudan in 2014 (site 9) and in Rwanda in 2013 (site 10) as described in ^7^.

Survey B: Samples were collected from the Nakuru (5 sites), Baringo (3), Machakos (2) and Kitui (2) regions of Kenya in August 2013. At the same time samples were collected from the Oromia region of Ethiopia (4).

### Initial Screening

Survey A: Total RNA was extracted using an RNeasy kit including the on - column DNase digestion (Qiagen, UK) and real time PCR for the presence of MCMV and SCMV was carried out as described in^5^.

Survey B: DAS - ELISA was performed on leaf samples for MCMV and indirect ELISA for SCMV as described in ^1^. Total RNA extraction was performed using Trizol (Ambion) according to manufacturer’s instructions.

### MiSeq Sequencing

For MiSeq sequencing indexed ScriptSeq V2 libraries were prepared using total RNA according to the manufacturer’s instructions (EpiBio, USA). Up to 20 different libraries were sequenced on a single 600 cycle V3 flow cell using an Illumina MiSeq. The resulting sequences were filtered to a quality score of 20 using Sickle ^33^ and assembled using Trinity ^34^. The resulting contigs were compared to the NCBI GenBank protein database (data download October 2015) using BLASTx + ^35^ and viral reads identified using MEGAN 5 ^36^. The number of reads mapping back to the viral genome was calculated using Bwa mem ^37^ and SamTools ^38^.

### HiSeq Sequencing

Indexed ScriptSeq V2 libraries were prepared using total RNA according to the manufacturer’s instructions (Epicentre). 30 libraries were sequenced on a single lane of an illumine HiSeq 2000 (BGI). Libraries had low quality bases trimmed using Trim galore! (threshold = 20)^39^ were assembled using Trinity^33^, then contigs compared to the NCBI Genbank nt database (October 2015) by BLASTn.

### MinION sequencing

For MinION sequencing the SQK - MAP005 2D cDNA kit (Oxford Nanopore) was used. All reagents were part of the kit unless alternative suppliers are detailed. Total RNA (250ng) was mixed with 0.5 μl of 250 μM random primers and 1 μl of 10 mM dNTP mix and the reaction volume adjusted to 13 μl with water. The reaction was incubated at 65 °C for 5 minutes then snap chilled on ice. First stand buffer (4 μl) and 2 μl 100 mM DTT were then added (both supplied with Superscript, Life Technologies) and the reaction incubated at 42 °C for 2 minutes. SuperScript II reverse transcriptase (1μl of 200U/μl - Life Technologies) was then added and the reaction incubated at 50 °C for 50 minutes followed by 70 °C for 15 minutes. Second strand synthesis reaction buffer (10μ) and 5 μl second strand synthesis enzyme mix from the second strand synthesis kit (NEB, UK) and 45 μl water were then added and the reaction incubated at 16 °C for 60 minutes. The resulting cDNA was then purified using 1.8 x AMPure XP beads (Agencourt) following the manufacturer’s instructions before being eluted in 52 μl water. The cDNA was end repaired and A tailed using the End Repair A - Tailing Kit (NEB, UK) following the manufacturer’s instructions before being purified using 1.8 x AMPure XP beads (Agencourt) following the manufacturer’s instructions before being eluted in 15 μl water.

PCR adapters (5 μl) and 20 μl Blunt/TA Ligase Master Mix (NEB, UK) were added to the cDNA and the mixture incubated at room temperature for 15 minutes. The adapted cDNA was then purified using 0.7 x AMPure XP beads (Agencourt) before being eluted in 25 μl water. The resulting cDNA was mixed with 2 μl amplification primers (provided in MAP005 kit), 50 μl LongAmp Taq 2x master mix (NEB) and 23 μl of water. The reaction was incubated at 95 °C for 3 minutes prior to 18 cycles of 95 °C for 15 seconds, 62 °C for 15 seconds and 65 °C for 10 minutes and a final incubation of 65 °C for 10 minutes on a C1000 PCR instrument (Biorad). The amplified cDNA was then purified using 0.7 x AMPure XP beads (Agencourt) before being eluted in 80 μl water.

A MinION sequencing library was prepared using the SQK - MAP005 2D DNA kit (Oxford Nanopore). The purified cDNA was end repaired and A - tailed using the End Repair A Tailing kit (NEB, UK) following the manufacturer’s instructions and again purified using 1.8 x AMPure XP beads (Agencourt) before being eluted in 31 μl water. Adapter mix (10 μl), 10 μl HP adapter and 10 μl Blunt/TA Ligase Master Mix (NEB, UK) were added to the cDNA and the mixture incubated at room temperature for 10 minutes. MyOne C1 Streptavidin beads (10 μl : Life Tech, UK) were washed twice in bead binding buffer, re - suspended in 100 μl bead binding buffer and mixed with the adapter ligated cDNA. After 5 min the mixture was placed on a magnetic stand, the supernatant removed. The beads were then washed twice by re - suspending in 200 μl bead binding buffer, replacing on the magnet and removing the supernatant. The beads were then re - suspended in 25 μl of elution buffer, incubated for 10 minutes at room temperature before replacing on the magnet and the purified library recovered. This library was called the “pre - sequencing mix”

The MinION (original MAP 2014 version) was connected to a Dell Latitude E7240 with MinKnow software (Oxford Nanopore) running. The Laptop was also running the Metrichor client (Oxford Nanopore) which transfers raw data from the MinION, uploads it to an instance running in the Amazon cloud and returns sequence data to the laptop. The R7.3 flowcell placed in the MinION was primed twice for 10 minutes with 150 μl prime mix (6.5 μl fuel mix, 162.5 μl running buffer and 156 μl water). The flow cell was then loaded with 150 μl library (6 μl pre - sequencing mix, 75 μl running buffer, 3 μl fuel mix, 66 μl water) and the 48 hr MAP script was then run. Poretools ^40^ was used to extract FASTA data from the MinKnow output. The resulting sequences were compared to the NCBI GenBank protein and nucleic acid database (Download date: October 2015) using BLASTx + ^35^ and viral reads identified using MEGAN 5 ^36^. The number of reads mapping to the viral genome was calculated using LAST ^41^ and SamTools ^38^. WIMP data was produced by re - uploading the raw MinION data using the Metrichor client (Oxford Nanopore).

Viral open reading frames and proteins were analysed using CLC mainworkbench (CLC Bio). Phylogenetic trees were constructed using MEGA 6 with 500 bootstrap resamplings ^42^.

## Acknowledgements

This work has received support through the Defra future proofing plant health project (PH0469), the Oxford Nanopore MAP programme, the Global Plant Clinic and the Swedish International Development Cooperation Agency (SIDA) through an award to the BecA - ILRI Hub

## Author contributions

IA, FS and LAB performed the experimental work and contributed to the preparation of the manuscript and figures. IA, NB, RM, LAB and JS wrote the main text of the manuscript. All authors reviewed the manuscript.

## Data availability statement

As stated in the text the sequences of the novel viruses are available on NCBI Genbank and the MinION data is available at the NCBI Short read archive.

